# Deformed Probability Estimation in Goal-Directed reinforcement learning model explains anxious-depression dimensions of psychiatric disorders

**DOI:** 10.1101/2025.07.16.665092

**Authors:** Sadjad Yazdani, Majid Nili Ahmadabadi, Babak Nadjar-Araabi, Abdol-Hossein Vahabie

**Affiliations:** School of Electrical and Computer Engineering - University of Tehran

**Keywords:** Computational psychiatry, Decision-Making Biases in Cognitive Modelling, Anxiety-Depression Dimension, Subjective Probability Distortion, Model-Based/Model-Free Learning Style, Description/Experience Gap

## Abstract

Psychiatric disorders are complex, multi-dimensional pathologies rooted in diverse cognitive processes. Computational psychiatry aims to reveal distortions in these processes through behavior modeling, providing a deeper understanding of psychiatric disorders. Previous studies, using Daw’s two-stage task, had linked the imbalance between habitual/model-free and goal-directed/model-based behaviors to disorders with compulsive behaviors and intrusive thoughts. The model-based component relies on the estimation of environmental probabilities. Therefore, we added a well-known deformation in subjective probability estimation to the model and this improved the model fitting. More importantly, the fitted deformation explains some variance of the anxious-depression dimension of psychiatric symptoms. The deformation parameter is aligned with the description-experience gap in decision-making literature. Our results point to subjective possibly distortion as the probable underlying cognitive process of anxiety, apathy, and depression. This study also shows that the inclusion of cognitive biases in modeling can extract the hidden aspects of behavior possibility linked to disorders. Our approach enhances the precision of computational psychiatry and provides deeper insights into the cognitive processes underlying psychiatric symptoms, paving the way for more effective, personalized therapeutic strategies.

**Significance Statement:** This study builds on a previously used model and Data that indicated a correlation between Model-Based preference and certain psychiatric disorders. By adding distortion to the probability estimation, we identified a new parameter correlated with depression and anxiety. The augmented model demonstrates an improved fit to behavior and aligns with Gillan’s previous findings. Our approach enhances the precision of computational psychiatry and provides deeper insights into the cognitive processes underlying psychiatric symptoms, paving the way for more effective, personalized therapeutic strategies.

## 1. Introduction

Nowadays, psychiatric disorders are a significant source of human discomfort in life, and researchers are very interested in understanding the underlying cognitive processes linked to these disorders. Several studies show that psychiatric disorders alter the brain’s information processing, leading to different behavior and cognition during different tasks [1, 2]. Computational psychiatry is an emerging field that seeks to understand these alterations by leveraging mathematical and computational tools to model the brain’s computational processes [3, 4]. Researchers in psychiatric modeling aim to uncover the underlying mechanisms of psychiatric symptoms based on behavioral indices [3, 5, 6, 7, 2]. Psychiatric modeling decomposes behavior into the underlying cognitive processes to have a better understanding of brain mechanisms. Research Domain Criteria (RDoC), as a modern categorizing framework, uses such models. The RDoC framework emphasizes understanding psychiatric disorders through dimensions of observable behavior and neurobiological measures, contrasting with the Diagnostic Statistical Manual (DSM), which classifies disorders based on symptom clusters and diagnostic criteria. To achieve these goals, various tasks and experimental paradigms have been employed such as the Stroop test, the Wisconsin Card Sorting Test, and various reinforcement learning tasks [4, 8, 9].

The reinforcement learning (RL) models extract the effect of expectation and reward on the choice behavior. The RL models reveal important aspects of behavior, from reward-seeking and decision-making to learning and plasticity. Researchers show that the dissociation of behavior’s building blocks, based on the RL models, has substantial explanatory and predictive power in many domains [10, 11, 12, 13]. The RL models associate habitual and goal-directed components of the behavior with Model-Free (MF) and Model-Based (MB) learning styles.

Nathaniel Daw and his colleagues explored this dissociation, particularly through their well-known two-stage task [14]. This task distinguishes the behavior between the MB approach, which involves planning and using a cognitive model of the environment, and the MF approach, which relies on habitual responses based on history of transitions and rewards. Gillan and her colleagues conducted a remarkable study analyzing the relationship between symptoms of various mental disorders and the balance between model-based (MB) and model-free (MF) learning styles, as extracted during Daw’s two-stage task. Their findings indicated that MB preference is associated with psychiatric symptoms aligned with a factor of compulsive behavior and intrusive thoughts [15]. However, there was no significant correlation with other disorders, particularly those aligned with the depression-anxiety factor.

The estimation of event probabilities has been extensively studied in various contexts, including anxiety and depression disorders. For instance, research has demonstrated that anxiety tends to increase the estimated probability of threatening events, while depression decreases the estimated probability of positive outcomes [16, 10, 12]. Deformation in probability estimation affects the MB component of RL through reward probabilities and transition probabilities, which represents the likelihood of moving from one state to another in the environment. This estimation process is crucial for planning and decision-making, as it enables individuals to predict the outcomes of their actions and select the most beneficial ones.

It is widely reported that people show a distorted subjective probability relative to the objective probability in decision-making situations. This distortion has been well-studied in prospect theory by Kahneman and Tversky [17, 18, 19]. Some studies support the idea that the probability distortion can be different in psychiatric disorders and related cognitive processes may play a role in the emergence of psychiatric disorders [12, 20].

In this study, we hypothesize that incorporating subjective probability deformation into the MB component of the RL model can capture possible distortions in probability estimation. This inclusion is expected to enhance the model fitting and to improve the predictive power of the model.

We incorporate the deformation of probability estimation to the model used by Gillan and her colleagues [15], as illustrated in Fig.1. Next, we fit our augmented model to the behavioral data, and extract the parameters, assuming a reliable range of parameters [21]. Obtained results indicate that the deformation parameter is strongly correlated with the depression-anxiety dimension of disorders.

**Fig 1.**
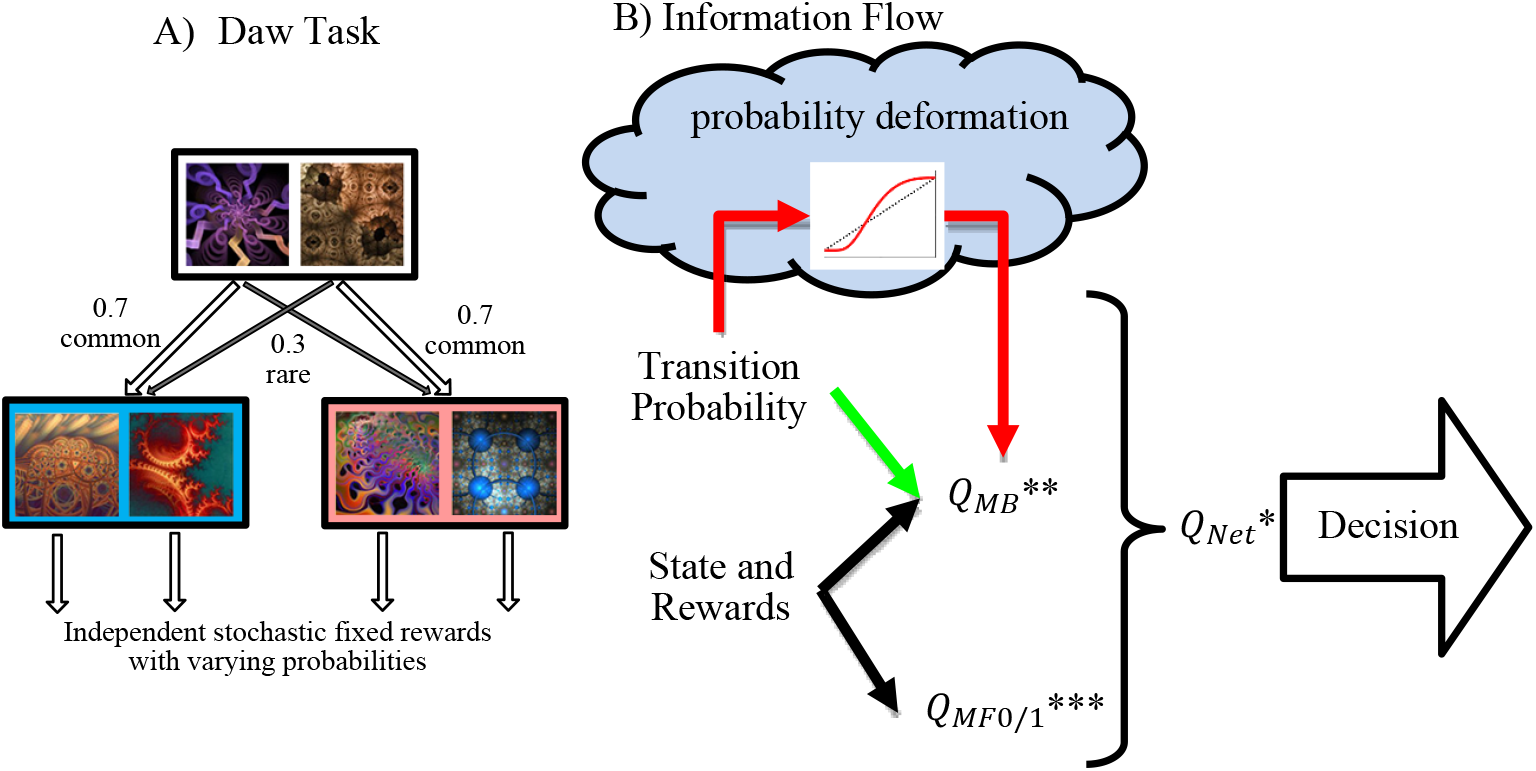
A) the Daw two-stage task, B) The flow of information in the computational model. The transition probability is used directly (green arrow) or it will deform (the red arrow) in the modeling. [* Eq(2), ** Eq(3) or Eq(5), *** SARSA-λ]

Consequently, our study demonstrates that integrating a relevant deformation component to the model can enhance the utility of the two-stage task in understanding compulsive disorders related to the anxiety-depression factor. This approach deepens our comprehension of brain function and psychiatric disorders.

## 2. Result

Model-based (MB) and Model-Free (MF) learning styles have been extensively utilized to investigate habitual and goal-directed models. The connection between MB approach preference and certain psychiatric disorders has been shown in the study of Gillan and her colleagues [15]. The MB approach relies on the transition probability of the environment. In this study, we hypothesized that different psychiatric disorders may differ in how this probability is estimated and applied in the model. Based on how participants handle transition probability in the model, as discussed in the method section, we compared five different models, summarized in Table 1.

**Table 1.**
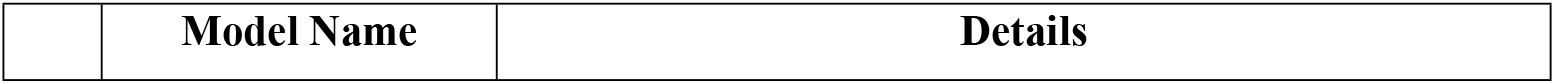
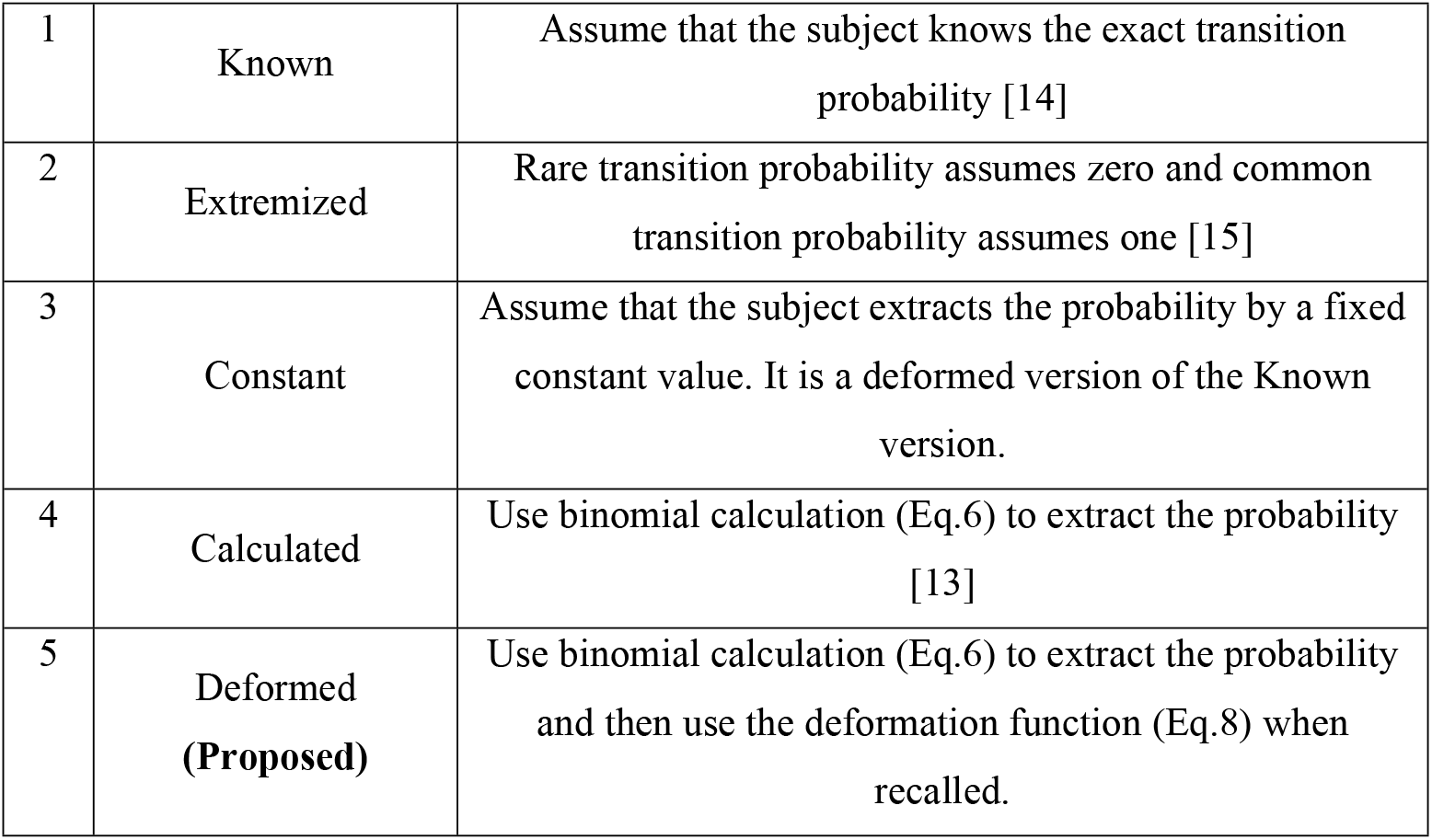
Models use different transition probabilities.

To evaluate models and to determine which model better predicts experimental data, we use the Akaike Information Criterion (AIC) per subject. As Table 2 shows, the model that used deformed probability gives us the best results.

**Table 2.** Model comparison. The average value of the Akaike Information Criterion per subject and the Standard Error are reported. There are 1413 data from Gillan’s study.

| Model | AIC average [SE] |
| --- | --- |
| Known | 282.41 [ 0.06] |
| Extremized | 282.39 [ 0.06] |
| Constant | 283.90 [ 0.06] |
| Calculated | 280.62 [ 0.06] |
| Deformed | **276.37 [ 0.08]** |

The winner model used the well-known Prelec one-parameter function [22] for the implementation of probability deformation. This function has a parameter *η* that determines how objective probability is transformed into subjective one. If *η* is one, there is no distortion, but if *η*<1 (*η*>*1)*, there is an overestimation (underestimation) of low probabilities and an underestimation (overestimation) of high probabilities; see Fig.2-A.

**Fig 2.**
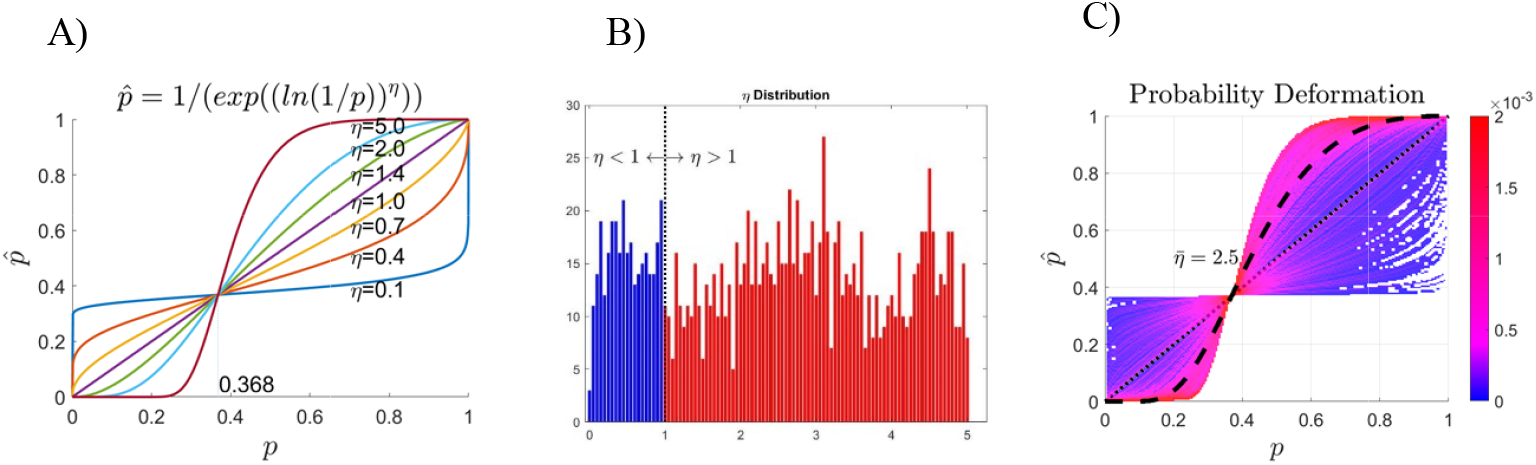
A) The deformation function (Prelec) for different η values. B) The distribution of per-subject fitted η. C) Distribution of deformation function based on model fitting; the average distribution is shown by a black line. The color code indicates the density, i.e. most subjects have *η* > 1.

Fig.2-B and Fig.2-C shows the distribution of fitted deformation parameter across the subjects. The average value of the fitted Prelec parameter is *η*=2.5. This indicates that the distribution of transition probabilities tends to underestimate low probabilities and overestimate high probabilities. This higher value is in contrast with studies that are description-based (for more detail, see the Discussion section). However, it is consistent with studies that are experience-based, and it is aligned with the reinforcement learning tasks [23, 24, 25].

The relationship between model parameters and psychiatric symptom indices reveals interesting information. In Table 3, the relationship between the total score of the self-questionary administered by Gillan et al. [15] is presented in relation to two model parameters: the subjects’ preference toward the MB approach (β*_M_*) and the amount of probability deformation (*η*). The first column of Table 3 replicates Gillan’s analysis on regression of one trial back. The other columns are regressions between parameters extracted from RL modeling and different symptom scores. Each row is the result of an independent analysis of one trial back regression or the regression of computational models in which the total score of the questionnaire related to each Symptom (z scored) or the factors are placed as *SymptomScorez* in the following linear model

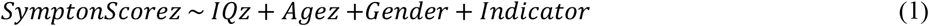

where the *IQz* and *Agez*, are z-scores of IQ and age, respectively. The Gender is either −1 or 1 for men and women, respectively. The indicator is either *β_MB_* (the MB weight) or *η* (the probability deformation parameter). The values presented in the table show the effect of the indicator on *SymptonScorez* from the regression analysis. For any subject, the preference toward MB style (*β_MB_*) was obtained by fitting both the extremized and the deformed computational model on its behavioral observations.

**Table 3.**
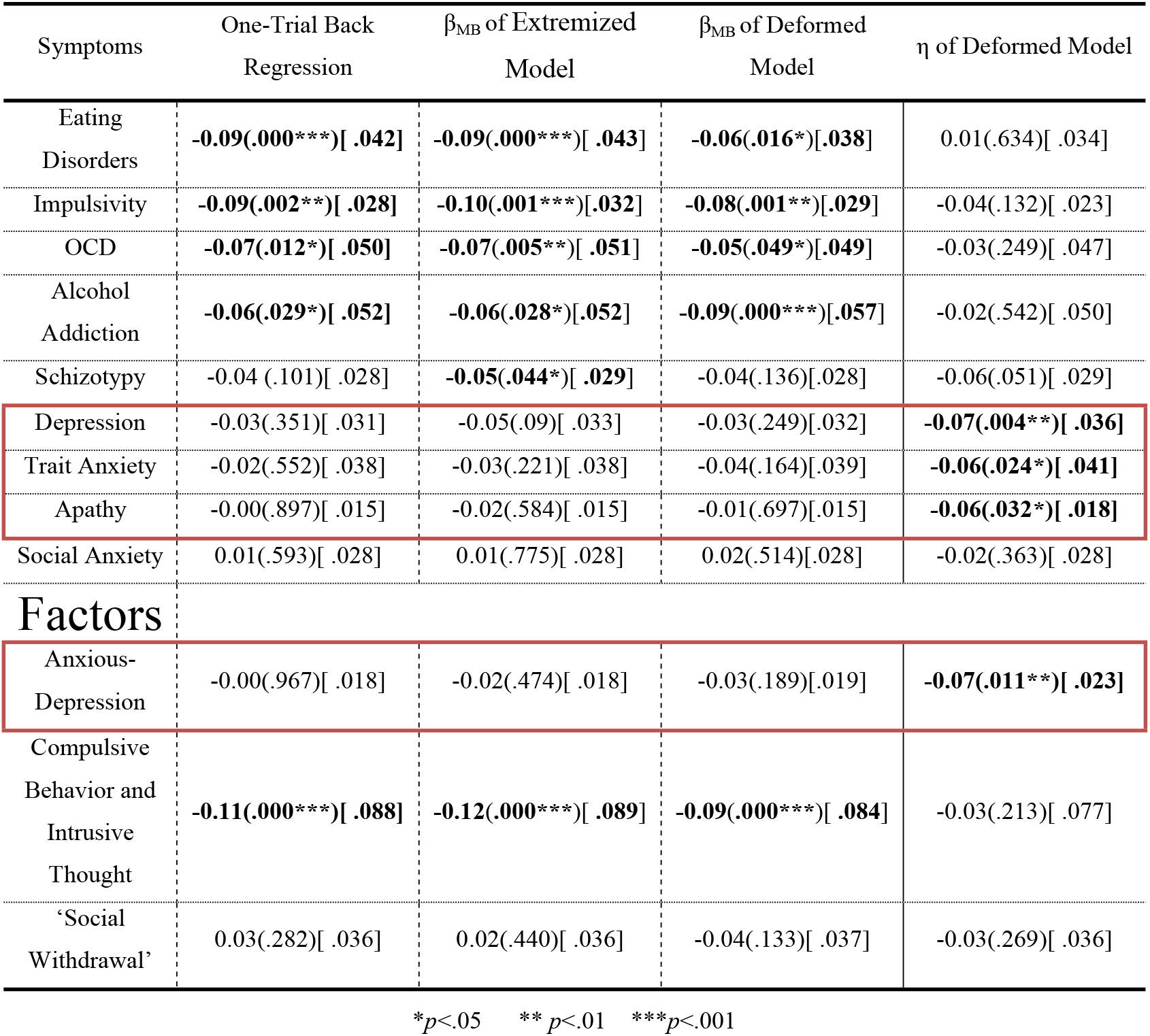
Comparing the predictive power of MB learning preference or subjective probability estimation deformation parameters using different computational models and one-trial back regression analysis. The one-trial back and Extremized models are replicated from the Gillan study [15]. Each box represents the effect of the indicator on symptom score in regression β (*p*-value) [*R^2^*]

| Symptoms | One-Trial Back<br>Regression | $\beta_{MB}$ of Extremized<br>Model | $\beta_{MB}$ of Deformed<br>Model | $\eta$ of Deformed Model |
| --- | --- | --- | --- | --- |
| Eating Disorders | **-0.09(.000\*\*\*)[.042]** | **-0.09(.000\*\*\*)[.043]** | **-0.06(.016\*)[.038]** | 0.01(.634)[.034] |
| Impulsivity | **-0.09(.002\*\*)[.028]** | **-0.10(.001\*\*\*)[.032]** | **-0.08(.001\*\*)[.029]** | -0.04(.132)[.023] |
| OCD | **-0.07(.012\*)[.050]** | **-0.07(.005\*\*)[.051]** | **-0.05(.049\*)[.049]** | -0.03(.249)[.047] |
| Alcohol Addiction | **-0.06(.029\*)[.052]** | **-0.06(.028\*)[.052]** | **-0.09(.000\*\*\*)[.057]** | -0.02(.542)[.050] |
| Schizotypy | -0.04 (.101)[.028] | **-0.05(.044\*)[.029]** | -0.04(.136)[.028] | -0.06(.051)[.029] |
| Depression | -0.03(.351)[.031] | -0.05(.09)[.033] | -0.03(.249)[.032] | **-0.07(.004\*\*)[.036]** |
| Trait Anxiety | -0.02(.552)[.038] | -0.03(.221)[.038] | -0.04(.164)[.039] | **-0.06(.024\*)[.041]** |
| Apathy | -0.00(.897)[.015] | -0.02(.584)[.015] | -0.01(.697)[.015] | **-0.06(.032\*)[.018]** |
| Social Anxiety | 0.01(.593)[.028] | 0.01(.775)[.028] | 0.02(.514)[.028] | -0.02(.363)[.028] |
| Factors |  |  |  |  |
| Anxious-Depression | -0.00(.967)[.018] | -0.02(.474)[.018] | -0.03(.189)[.019] | **-0.07(.011\*\*)[.023]** |
| Compulsive Behavior and Intrusive Thought | **-0.11(.000\*\*\*)[.088]** | **-0.12(.000\*\*\*)[.089]** | **-0.09(.000\*\*\*)[.084]** | -0.03(.213)[.077] |
| 'Social Withdrawal' | 0.03(.282)[.036] | 0.02(.440)[.036] | -0.04(.133)[.037] | -0.03(.269)[.036] |

\* $p < .05$ \*\* $p < .01$ \*\*\* $p < .001$

According to Table 3, Eating Disorder, Impulsivity, OCD, and Alcohol Addiction are correlated with the subject preference towards the MB approach, based on behavioral regression analysis and both computational models. In addition, Schizotypy has a significant relationship with MB preference, based on the model provided by Gillan and her colleagues, while this relationship in the proposed model and regression analysis is not meaningful. In other words, the proposed model is more consistent with behavioral regression.

On the other hand, the probability estimation deformation parameter has a significant relationship with some symptoms not explained by the Gillan model. Based on the results in Table 3, the amount of subjective probability deformation (*η*) has a meaningful relation to Depression, Trait Anxiety, and Apathy indices. These disorders account for the most part of Anxious-Depression factor based on the Gillan factor analysis [15]. Consequently, factor 1 is significantly related to the probability estimation deformation parameter *η*, as well.

## 3. Discussion

In a previous study, Gillan et al. demonstrated that the learning styles, Model-Based (MB) and Model-Free (MF) behavior, is correlated with some psychiatric symptoms associated with compulsive behavior. The MB approach requires calculating the state transition probability. Here, we implemented a new model incorporating deformations in subjective probability estimation. Proposed augmented model confirmed the explained variance of compulsive behaviors by MB and MF behaviors, reported in [15] and [26]. On top of that, our model showed the capability of explaining some other variance in the anxiety-depression dimension of psychiatric symptoms using estimated subjective probability distortion. The proposed model exhibited superior fittings compared to previous models. Moreover, the distortion in subjective probability is aligned with the description-experience gap in the literature and, showing underestimation and overestimation of low and high probabilities, respectively, as opposed to description-based distortions.

An effective approach in learning models involves integrating well-known cognitive biases into the model to improve behavior predictability. For example, Palminteri et al demonstrated that incorporating confirmation bias into model can improve fitting accuracy [27]. Daniel Kahneman and Amos Tversky studied various cognitive biases [17], which often influence behavior unconsciously. One notable bias that introduced in prospect theory literature is the distortion in subjective probability [17]. On the other hand, the MB learning style relies on the estimation of state transition probability and typically deformation has been ignored in modeling MB learning. In our proposed method, we incorporated a subjective probability estimation parameter in the model. Our results indicate that the subjective probability bias is linked to the anxiety-depression dimension and accounts for some variance of depression, trait anxiety, and apathy.

These findings align well with the existing literature on anxiety and depression. Probability bias is a typical example of a cognitive distortion linked to anxiety disorders [16, 28]. Biased probability estimates of emotional events in anxious individuals are usually explained in terms of ‘‘availability heuristic’’, which suggests that individuals estimate the likelihood of an event not through logical calculation but based on the ease with which similar scenarios can be retrieved from memory [18]. Probability distortion is also observed in individuals with depression [29]. Hagiwara et al. investigated nonlinear probability weighting in depression and anxiety [12]. Pulcu and his colleague explored how individuals with major depressive disorder (MDD) and generalized anxiety disorder (GAD) perceive probabilities and make decisions under risk [30]. Individuals with anxiety disorders often exhibit heightened sensitivity to potential threats and may overestimate the likelihood of negative outcomes and people with depression may exhibit a negative bias in their learning from experience, leading to an underestimation of positive outcomes and an overemphasis on negative feedback. Our study shows that subjective probability estimation plays a role in depression-anxiety dimension. Moreover, schizotypy showed a near-significant relationship with subjective probability distortion. It can be associated with paranoid thoughts and help to understand why these people have a negative prediction about others’ intentions.

Most studies on subjective probability are description-based and demonstrate an overestimation of low probabilities and an underestimation of high probabilities. However, our fitted model is related to experience-based probability estimation. The fitted Prelect parameter was greater than one (η >1) for most subjects, indicating an underestimation of low probabilities and an overestimation of high probabilities. This pattern is known as S-shape and is inverse pattern relative to description-based average behavior. Literature highlights a difference between description-based and experience-based probability estimation. The description-experience gap refers to the phenomenon where individuals make different decisions based on whether they receive information through descriptive statistics (e.g., probabilities presented numerically) or through direct experience (e.g., probability estimation based on personal observations). This gap has been investigated in several studies [31, 32, 33, 34] and their findings are consistent with our fitted parameters. Our study provides strongest evidence (based on participant number) and approval for the S-shape pattern of probability estimation deformation in experience-based estimation. This finding is particularly relevant in the context of psychiatric disorders and suggests that there is a clear difference in the prediction of upcoming events based on the experience, compared to description cases, in anxious and depressed individuals.

In line with the literature, we did not find any significant relation between probability deformation and impulsivity, OCD, alcohol addiction, and eating disorders indices although these symptoms were related to goal-directed versus habitual trade-off. In the study by Gillan et al, a factor analysis of psychiatric symptoms identified three main dimensions [15]. Our results show that with precise modeling of cognitive processing, Daw’s two-stage task is effective in explaining some variance of two main dimensions. The third dimension is related to social aspects and because the task is not developed in a social context, it is expected that we did not find any explained variance there. Future studies are needed to design a social version of this task and link the parameter in the social version to the third dimension [35].

Finally, we know that there are multiple approaches to categorizing psychiatric disorders. The DSM (Diagnostic Statistical Manual) defines psychiatric disorders based on traditional classifications and overwhelming clinical symptoms, while RDoC (Research Domain Criteria) employs an increasingly embraced multidimensional approach based on biological and cognitive sciences in understanding psychiatric disorders. The present study demonstrated how MB and MF behaviors and deformations in subjective probability estimation can be linked to various dimensions of psychiatric symptoms. We modeled the underlying cognitive processes during the well-known Daw two-stage task and extracted two parameters that relate to different psychiatric symptom dimensions. This parallel separation of psychiatric dimensions with cognitive process decomposition exemplifies what the RDoC framework aims to capture in psychiatric domains. Such research contributes toward further enriching the RDoC framework and enabling a more efficient understanding and treatment of psychiatric disorders.

As shown in our study, it is more effective to extract the relevant parameters in experience-based methods, which is only possible through behavioral modeling because such parameters clarify the cognitive processes that are underlying layers of human behavior. So, computational psychiatry and RDoC can be considered complementary approaches to understanding psychiatric disorders. Our research underscores the importance of knowledge-informed computational elements in computational psychiatry. It also highlights the need for more profound computational models to address the complexities encountered in psychiatric disorders and to better understand behavior.

Understanding these computational processes may allow researchers to learn more about how various aspects of cognition and decision-making are affected by mental health phenomena. The inclusion of components such as probability deformation, closely creates an ecologically valid model, wherein subjects’ perceptions of probabilities differ from the actual probabilities. This difference could significantly affect the way such individuals would make decisions and, more importantly, their mental health condition. Addressing such distortions is the key to developing new and enriched intervention strategies and therapeutic modalities. Computational models help in the measurement and analysis of these barely noticeable but potent discrepancies in human behavior. With this approach, treatments can be better personalized into precision medicine for effectiveness in mental health care.

## 4. Method

Daw and his colleagues present a computational model that combines the values of the choices based on model-based (MB) and model-free (MF) systems linearly [14]. In 2013, Otto and his colleagues reparametrized the model presented by Daw and his colleagues [36]. Gillan and her colleagues studied the relationship between psychiatric disorders and subjects’ preference for the MB approach using regression analysis of behaviors and a modified version of the Otto et al.’s model [15]. In this modified model, the value of each choice is obtained by a linear combination of MB and two MF systems according to the following equation:

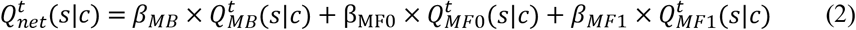

In this equation, 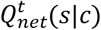 represents the final value of the choice *c* in the *t*th attempt in the state *s*. This value is obtained by a linear combination of three components: the MB value *Q_MB_*, and two MF values *Q*_*MF*0_ and *Q*_*MF*1_, calculated using two SARSA-λ algorithms with λ equal to zero (*MF*0) and one (*MF*1), respectively.

In the MB component, valuation is carried out through various calculations, and the probability of transition between states influences the evaluation of each future choice. Researchers have employed different approaches to handle the transition probability value including assuming a fixed known parameter or using a numerical method to calculate it.

Gillan and her colleagues considered the value of *Q_MB_* to be equal to the transactions received in the second stage, assuming the dominant transfer [15].

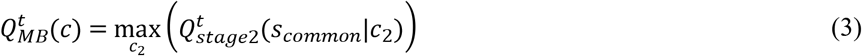

We refer this method as “extremized” due to the use of the extreme value of probability transactions. The value of 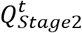 is updated according to the delta rule with the reward *r_t_*and the learning rate *α* :

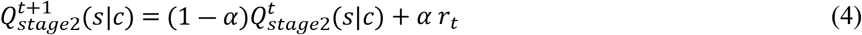

In the model presented by Daw and his colleagues, the values of MB in the first stage are calculated in line with the Bellman equation, considering the transition probability [14].

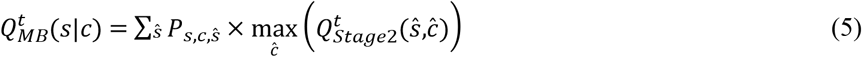

Where *P*_*s,c,ŝ*_ is the probability of transition from state *s* to state *s* during choice *c*. In the Daw task, this probability is fixed and equal to 0.7 and 0.3, respectively, for common and rare transfers, although the subject does not have accurate information about it and catches this information during the performing task or by pre-train process.

Daw and his colleagues assumed in their study that subjects extracted the transition probabilities in pre-training and the first few trials, subsequently using their accurate values (0.7 and 0.3) for MB calculations [14]. We refer to this method as “known.” Also, it seems worthwhile to assume that the participants perceive the transition probability for common as a fixed constant value, with the rare transition probability derived from this constant. We label this method as “constant.” On the other hand, for an environment with probabilistic transitions between states and a constant transition probability, it can be calculated numerically in general.

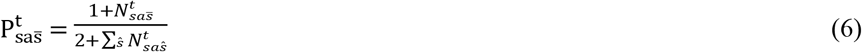

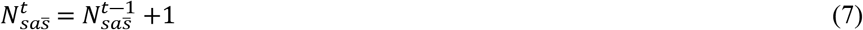

This method is referred as “calculated”. Finally, in the proposed method (deformed), the transition probability distorted through a deformation function after calculation.

Researchers model the probability distortion in the human mind using different methods [22]. A widely used function for this purpose is the one-parameter Prelec function. In this study, we consider Prelec for this deformation function as follows:

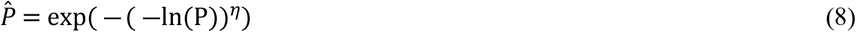

A suitable value of *η* as a free parameter of this function can create the desired deformation. In this study, we performed this modeling and extracted the parameters by fitting the model using maximum likelihood estimation in a reliable range of parameters. [21]. The Prelec parameter (η) was assumed to range from zero to five. It should be noted that a larger value for η does not change the deformation effectively and the results remain stable with larger ranges.

## Data and Code availability

The data that is used in this study is available online by Gillan on OSF database [37].

Also, all codes are available on GitHub [38].

## Acknowledgment

We would like to express our gratitude to Clare Gillan for sharing the data. The authors declare that no grants, financial support, or material resources were received for the conduct, authorship, or publication of this research.

## Disclosures

The authors declare no competing financial interests or other conflicts of interest.

## Notes

### Competing Interest Statement

The authors have declared no competing interest.

## References

[1] S. Kelly, S. Guimond, A. Lyall, W. S. Stone, M. E. Shenton, M. Keshavan and L. J. Seidman, ‘Neural correlates of cognitive deficits across developmental phases of schizophrenia,’ Neurobiology of Disease, vol. 131, p. 104353, 2019.

[2] A. Talwar, ‘Computational Models Describe Individual Differences in Cognitive Function and Their Relationships to Mental Health Symptoms,’ UCL, 2023.

[3] A. Anticevic and J. D. Murray, Computational Psychiatry: Mathematical Modeling of Mental Illness, Academic Press, 2017.

[4] R. A. Adams, Q. J. M. Huys and J. P. Roiser, ‘Computational Psychiatry: towards a mathematically informed understanding of mental illness,’ J Neurol Neurosurg Psychiatry, vol. 87, no. 1, pp. 53–63, 2016.

[5] J. Gläscher, N. Daw, P. Dayan and J. P. O’Doherty, ‘States versus Rewards: Dissociable Neural Prediction Error Signals Underlying Model-Based and Model-Free Reinforcement Learning,’ Neuron, vol. 66, no. 4, p. 585–595, 2010.

[6] Q. J. M. Huys, T. V. Maia and M. J. Frank, ‘Computational psychiatry as a bridge from neuroscience to clinical applications,’ Nature Neuroscience, vol. 19, no. 3, p. 404–413, 2016.

[7] P. Karvelis, M. P. Paulus and A. O. Diaconescu, ‘Individual differences in computational psychiatry: A review of current challenges,’ Neuroscience & Biobehavioral Reviews, vol. 148, p. 105137, 2023.

[8] C. Hedge, A. Bompas and P. Sumner, ‘Task Reliability Considerations in Computational Psychiatry,’ Biological Psychiatry: Cognitive Neuroscience and Neuroimaging, vol. 5, no. 9, pp. 837–839, September 2020.

[9] A. Steinke, F. Lange, C. Seer and B. Kopp, ‘Towards a Computational Cognitive Neuropsychology of Wisconsin Card Sorts: A Showcase Study in Parkinson’s Disease,’ Computational Brain & Behavior, p. 137–150, 2018.

[10] R. W. Booth and D. Sharma, ‘Biased probability estimates in trait anxiety and trait depression are unrelated to biased availability.,’ Journal of Behavior Therapy and Experimental Psychiatry, vol. 73, no. 27, p. 101672, 2021.

[11] C. M. Gillan and T. W. Robbins, ‘Goal-directed learning and obsessive-compulsive disorder,’ Philosophical Transactions of the Royal Society B, vol. 369, no. 1655, p. 20130475, 2014.

[12] K. Hagiwara, Y. Mochizuk, C. Chen, H. Lei, M. Hirotsu, T. Matsubara and S. Nakagawa, ‘Nonlinear Probability Weighting in Depression and Anxiety: Insights From Healthy Young Adults,’ Front. Psychiatry, vol. 13, p. 810867, 2022.

[13] A. Konovalov and I. Krajbich, ‘Gaze data reveal distinct choice processes underlying model-based and model-free reinforcement learning,’ Nature Communications, vol. 7, p. 12438, 2016.

[14] N. D. Daw, S. J. Gershman, B. Seymour, P. Dayan and R. J. Dolan, ‘Model-based influences on humans’ choices and striatal prediction errors,’ Neuron, vol. 69, no. 6, p. 1204–1215, 2011.

[15] C. M. Gillan, M. Kosinski, R. Whelan, E. A. Phelps and N. D. Daw, ‘Characterizing a psychiatric symptom dimension related to deficits in goal-directed control,’ eLife, vol. 5, p. e11305, 2016.

[16] S. j. Bishop and C. Gagne, ‘Anxiety, Depression, and Decision Making: A Computational Perspective,’ Annual Review of Neuroscience, vol. 41, no. 1, p. 371–388, 2018.

[17] D. Kahneman and A. Tversky, ‘Kahneman, D., & Tversky, A. (). Prospect Theory: An Analysis of Decision under Risk,’ Econometrica, vol. 47, no. 2, p. 263–292, 1979.

[18] A. Tversky and D. Kahneman, ‘Judgment under Uncertainty: Heuristics and Biases.,’ Science, vol. 185, no. 4157, p. 1124–1131, 1974.

[19] A. Tversky and D. Kahneman, ‘Advances in prospect theory: Cumulative representation of uncertainty,’ Journal of Risk and Uncertainty, vol. 5, no. 4, p. 297–323, 1992.

[20] Q. J. Huys, J. T. Vogelstein and P. Dayan, ‘Psychiatry: Insights into depression through normative decision-making models,’ Neuroscience, vol. 21, pp. 1–8, 2009.

[21] S. Yazdani, A.-H. Vahabie, B. Nadjar-Araabi and M. Nili Ahmadabadi, ‘Better than maximum likelihood estimation of model-based and model-free learning style.,’ Basic and Clinical Neuroscience (BCN), 2025.

[22] P. Liu, A. Schied and R. Wang, ‘Distributional Transforms, Probability Distortions, and Their Applications,’ SSRN Electronic Journal, 2019.

[23] S. Ferrari-Toniolo, P. M. Bujold and W. Schultz, ‘Probability distortion depends on choice sequence in rhesus monkeys,’ Journal of Neuroscience, vol. 39, no. 15, p. 2915–2929, 2019.

[24] B. Garcia, F. Cerrotti and S. Palminteri, ‘The description-experience gap: A challenge for the neuroeconomics of decision-making under uncertainty,’ Philosophical Transactions of the Royal Society B, vol. 376, no. 1819, 2021.

[25] W. R. Stauffer, A. Lak, P. Bossaerts and W. Schultz, ‘Economic choices reveal probability distortion in macaque monkeys,’ Journal of Neuroscience, vol. 35, no. 7, p. 3146–3154, 2015.

[26] C. M. Gillan, E. Kalanthroff, M. Evans, H. M. Weingarden, R. J. Jacoby, M. Gershkovich, I. Snorrason, R. Campeas, C. Cervoni, N. C. Crimarco, Y. Sokol, S. L. Garnaat, N. C. R. McLaughlin, E. A. Phelps, A. Pinto, C. L. Boisseau, S. Wilhelm, N. D. Daw and H. B. Simpson, ‘Comparison of the Association Between Goal-Directed Planning and Self-reported Compulsivity vs Obsessive-Compulsive Disorder Diagnosis,’ JAMA Psychiatry, vol. 77, no. 1, pp. 77–85, 2020.

[27] S. Palminteri, G. Lefebvre, E. J. Kilford and S.-J. Blakemore, ‘Confirmation bias in human reinforcement learning: Evidence from counterfactual feedback processing., 13(8,’ PLoS Computational Biology, vol. 13, no. 8, p. e1005684, 2017.

[28] G. Butler and A. Mathews, ‘Cognitive processes in anxiety,’ Advances in Behaviour Research and Therapy, vol. 5, no. 1, p. 51–62, 1983.

[29] T. Kube, R. Schwarting, L. Rozenkrantz, J. A. Glombiewski and W. Rief, ‘Distorted Cognitive Processes in Major Depression: A Predictive Processing Perspective,’ Biological Psychiatry, vol. 87, no. 5, pp. 388–398, 2020.

[30] E. Pulcu, P. D. Trotter, E. J. Thomas, M. McFarquhar, G. Juhasz, B. J. Sahakian, J. F. W. Deakin, R. Zahn, I. M. Anderson and R. Elliott, ‘Temporal discounting in major depressive disorder,’ Psychological Medicine, vol. 44, no. 9, pp. 1825–1834, 2014.

[31] M. Abdellaoui, O. L’Haridon and Corina Paraschiv, ‘Experienced vs. Described Uncertainty: Do We Need Two Prospect Theory Specifications?,’ Management Science, vol. 57, no. 10, pp. 1879–1895, 2011.

[32] I. Aydogan and Y. Gao, ‘Experience and rationality under risk: re-examining the impact of sampling experience,’ Experimental Economics, vol. 23, p. 1100–1128, 2020.

[33] R. Cubitt, O. Kopsacheilis and C. Starmer, ‘An inquiry into the nature and causes of the Description - Experience gap.,’ Journal of Risk and Uncertainty, vol. 65, no. 2, p. 105–137, 2022.

[34] R. Hau, T. J. Pleskac, J. Kiefer and R. Hertwig, ‘The description-experience gap in risky choice: The role of sample size and experienced probabilities,’ Journal of Behavioral Decision Making, vol. 21, no. 5, p. 493–518, 2008.

[35] P. L. Lockwood, M. Klein-Flügge, A. Abdurahman and M. J. Crockett, ‘Neural signatures of model-free learning when avoiding harm to self and other,’ bioRxiv preprint, 2019.

[36] A. R. Otto, C. M. Raio, A. Chiang, E. A. Phelps and N. D. Daw, ‘Working-memory capacity protects model-based learning from stress,’ Proceedings of the National Academy of Sciences, vol. 110, no. 52, p. 20941–20946, 2013.

[37] C. Gillan, ‘Characterizing a psychiatric symptom dimension related to deficits in goal-directed control,’ 09 06 2021. [Online]. Available: https://osf.io/usdgt/.

[38] S. Yazdani, ‘sadjad-yazdani/DeformationPaper,’ 2025. [Online]. Available: https://github.com/sadjad-yazdani/DeformationPaper.

